# Different Physical Parameters in the Bonds between Platelet Glycoprotein Ibα with von Willebrand factor and with Coagulant Factor XI -Results from the Molecular Dynamic Simulation-

**DOI:** 10.1101/2023.04.16.534191

**Authors:** Masamitsu Nakayama, Shinichi Goto, Satoko Takemoto, Hideki Oka, Hideo Yokota, Shu Takagi, Shinya Goto

**Affiliations:** Department of Medicine (Cardiology), Tokai University School of Medicine, Isehara, Japan; Division of General Internal Medicine & Family Medicine, Department of General and Acute Medicine, Tokai University School of Medicine, Isehara, Japan; Division of Cardiovascular Medicine, Department of Medicine, Brigham and Women’s Hospital, Boston, USA; Image Processing Research Team, Center for Advanced Photonics, RIKEN, Wako, Japan; Graduate School of Engineering, The University of Tokyo, Tokyo, Japan

**Keywords:** Platelet, von Willebrand factor, glycoprotein Ibα, coagulation factor XI, molecular dynamics

## Abstract

**Background:** Both von Willebrand factor (VWF) and coagulation factor XI (FXI) bind with platelet membrane glycoprotein (GP) Ibα. However, the differences in the physical parameters in the bonds between VWF-GPIbα and FXI-GPIbα mediating different biological functions are unclear.

**Methods:** The FXI molecule was arranged in 9 different initial positions around the structure of GPIbα bound to VWF. The position coordinate and velocity vectors of all atoms constructing VWF, GPIbα, and FXI were calculated in each 2 femto (10^−15^) second using the Chemistry at HARvard Macromolecular Mechanics (CHARMM) force field. The physical parameters of VWF-GPIbα and FXI-GPIbα bonds were calculated by molecular dynamic (MD) simulations.

**Results:** MD calculation revealed 2.8 to 11.3 times greater positional fluctuations in atoms constructing FXI-GPIbα as compared to those constructing VWF-GPIbα (RMSDs: 5.9±1.5 to 18.1±7.9 Å for FXI-GPIbα vs 1.6±0.1 to 2.1±0.3 Å for VWF-GPIbα). The absolute value of non-covalent binding energy generated in FXI-GPIbα (65.5±79.7 to 517.6±54.2 kcal/mol) was smaller than that generated in VWF-GPIbα (678.5±58.3 to 1000.4±75.1 kcal/mol). The binding structure of VWF-GPIbα was stable and was only minimally influenced by the presence of FXI-GPIbα binding.

**Conclusions:** Our MD calculation results revealed that atoms constructing the VWF-GPIbα bond are physically more stable and produce more non-covalent binding energy than the bond of FXI-GPIbα. The physical parameters in the VWF-GPIbα bond were not largely influenced by FXI binding with GPIbα.

---

Platelets play crucial roles in regulating vascular functions.^1–4^ They accumulate promptly at vessel injury sites to stop the bleeding. ^2, 5, 6^ Under high wall shear rate conditions, initial adhesion of platelets at vessel injury sites is mediated exclusively by their membrane glycoprotein (GP) Ibα binding with the A1 domain of von Willebrand factor (VWF) expressed at the site of the injury. ^7–11^

Stable bindings of GPIbα to the VWF could only be detected in specific conditions such as in the presence of ristocetin^12^, botorocetin^13^ or with the “gain of function” mutations in either in the GPIbα or in the VWF. ^14–17^ Without them, the binding of GPIbα with VWF is transient. ^8^ The apparently stable binding of platelets to VWF was detected after exposure to high shear stress, but this stable binding was mediated by the binding between VWF and focally/segmentally activated GPIIb/IIIa on platelets that were initially transiently bound to VWF. ^18^ Accordingly, the VWF-GPIbα bond generates strong binding force transiently to capture flowing platelet on the injured vessel wall. The underlying physical mechanism of the unique function of the VWF-GPIbα bond as compared to other receptor-ligand interactions has yet to be clarified.

Karplus et al. developed a force field with coarse-grained quantum mechanics (QM) integrated into molecular mechanics (MM). Their technology, namely Chemistry at Harvard Macromolecular Mechanics (CHARMM), enables the prediction of the dynamic three-dimensional structures and physical parameters of various macromolecules under various conditions with computer-based molecular dynamic (MD) simulation. ^19, 20^ The energetically stable binding structure of wild-type GPIbα bound to the wild-type VWF was calculated by MD simulation. ^21^ The validity of the calculated structure-based physical parameters ^21^ was confirmed mechanically by comparing the predicted binding force between the two molecules with the experimental measurements using atomic force microscopy ^22^ and optical tweezers.^23^ Moreover, the validity of calculation results were also confirmed by the prediction of lower binding energy between the loss-of-function single point mutation (G233D) of GPIbα with VWF. ^24, 25^ The MD simulation could be applied to identify the specific physical characteristics of the VWF-GPIbα bond in comparison with other protein bonds such as GPIbα binding with coagulant proteins. The platelet GPIbα is known to bind with various coagulant proteins such as thrombin, and coagulation factor XI (FXI). ^26–28^ These coagulant proteins binding with GPIbα enhance the local activation of coagulation cascade around platelets but neither contribute to platelet adhesion nor aggregation. ^26, 29^

Here, potentially different physical characteristics of the same receptor protein of GPIbα binding with different ligands of VWF (VWF-GPIbα bond) and FXI (FXI-GPIbα bond) were investigated by molecular dynamic simulation calculations.**Methods**.

## 1. Initial Structure for MD simulation

The previously published stable structure of the A1 domain of the wild-type von Willebrand factor (VWF: residues ASP(D):1269-PRO(P):1472) binding with the N-terminal domain of platelet glycoprotein Ibα (GPIbα: residues HIS(H):1-PRO(P):265) was used as the initial binding structure of GPIbα-VWF bond. ^21^ Three-dimensional structure of FXI solved by X-ray crystallography was used as the initial structure of FXI. (PDBID: 2F83^30^) Previous research has shown that the GPIbα binding sites within FXI were constructed of positively charged amino acids (K252 and K253). ^26, 31^ However, the FXI-binding sites within GPIbα are unknown. Thus, the negatively charged amino acids (D and E) appearing on the GPIbα molecule(s) shown in **Fig. 1** were identified as the potential binding site(s) for FXI. The 15 negatively charged amino acids within GPIbα were grouped in 5 sets of positions, that potentially interact with K252 and K253 in FXI based on the 3D coordinates: D252-D249, E181-D159-E135, E66-D21, 172-E125 and D73-D28. (**Fig. 1**) FXI was placed at 9 different positions as shown by **Supplemental pdb files** (A.pdb to I.pdb) so that the K252/K253 region was close enough to interact with these 5 positions.

**Fig. 1.**
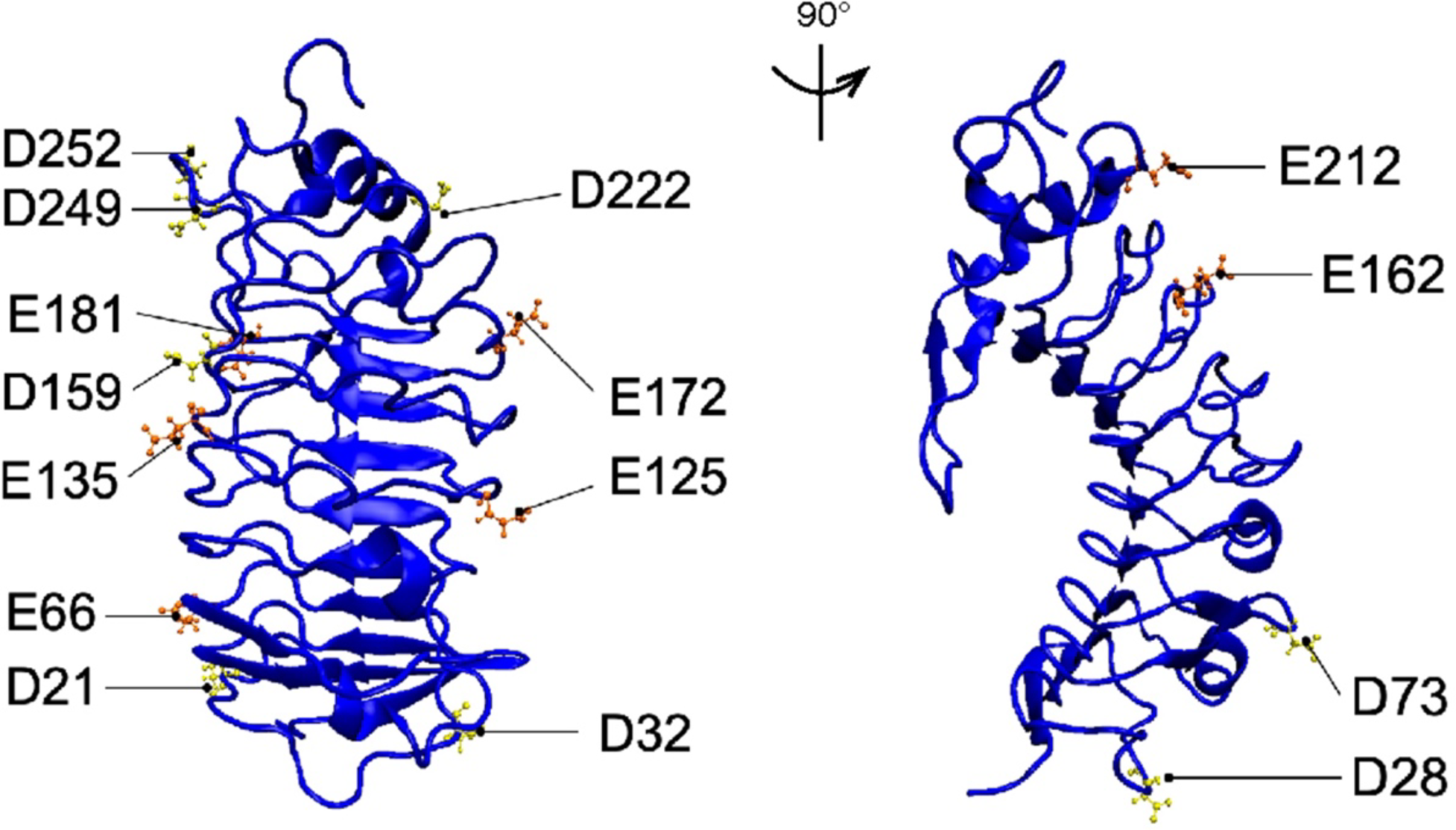
The Negatively Charged Amino Acid on GPIbα. Negatively charged amino acids of aspartic acid (D) and glutamic acid (E) in GPIbα are shown as yellow and orange Corey-Pauling-Koltun (CPK) model. The left panel shows the views of GPIbα from the opposite side of the VWF binding site and the right panel shows views after a 90-degree clockwise rotation around the vertical axis from that position. The amino acids of D252/D249, E181/D159/E135, E66/D21, E172/E125, and D73/D28 were considered as the groups of amino acids interacting with K252/K253 in FXI based on the position in the 3D coordinates.

## 2. MD Simulation Calculation

From all 9 initial positions of FXI arranged around GPIbα, Newton’s second law (shown below) was numerically solved for all atoms constructing the defined VWF binding region of GPIbα (HIS(H):1-PRO(P):265), GPIb binding site of VWF (residues ASP(D):1269-PRO(P):1472), and FXI (whole molecule) along with the surrounding water molecules with the Nanoscale Molecular Dynamics (NAMD) software package.^32^

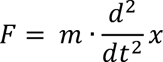

Where *F*, *m*, and *x* denote the force, mass, and displacement, respectively.

The simulation was performed on computers equipped with 4 sets of NVIDIA® Tesla® V100 GPU (HPC5000-XSLGPU4TS, HPC systems Inc., Tokyo, Japan). As the force field, CHARMM-36 was used for all simulations. The water molecules were modeled as CHARMM TIP3P (transferable intermolecular potential with three interaction sites).^33^ The position coordinates and velocity vectors of each atom and water molecule were calculated in each 2.0 femtosecond (2 × 10^−15^ s). A particle mesh Ewald (PME) treatment for long-range electrostatics was applied. ^34, 35^ The cut-off length of 12 Å was set as the maximum distance allowing direct interactions. Visual Molecular Dynamics (VMD) version 1.9.3 ^36^ was used for visualization of the results.

Of the huge calculation results obtained from 450 ns, snapshots were selected for every 10 ns. A total of 45 images each was selected from the calculation results starting from the nine different initial conditions and were used to compile **Supplemental Movies 1**. Since calculation results stabilized at approximately 250 ns, the results obtained after 400 ns (400 - 450 ns) were used to analyze the physical parameters such as the root mean square deviations (RMSDs), non-covalent binding energy, and the salt-bridges. A total of 50 images selected in every 1 ns within 400 to 450 ns were used to compile **Supplemental Movie 2**. For obtaining the results shown in **Fig. 2b**, the snapshots obtained every 1 ns from 400 to 450 ns were converted into 256 scale color images using ImageJ software (NIH, USA). Then, the 50 images obtained from different initial positions of FXI were overlaid with an alpha value of 0.02 to construct panel b in **Fig. 2** according to the previous publication. ^37^

**Fig. 2.**
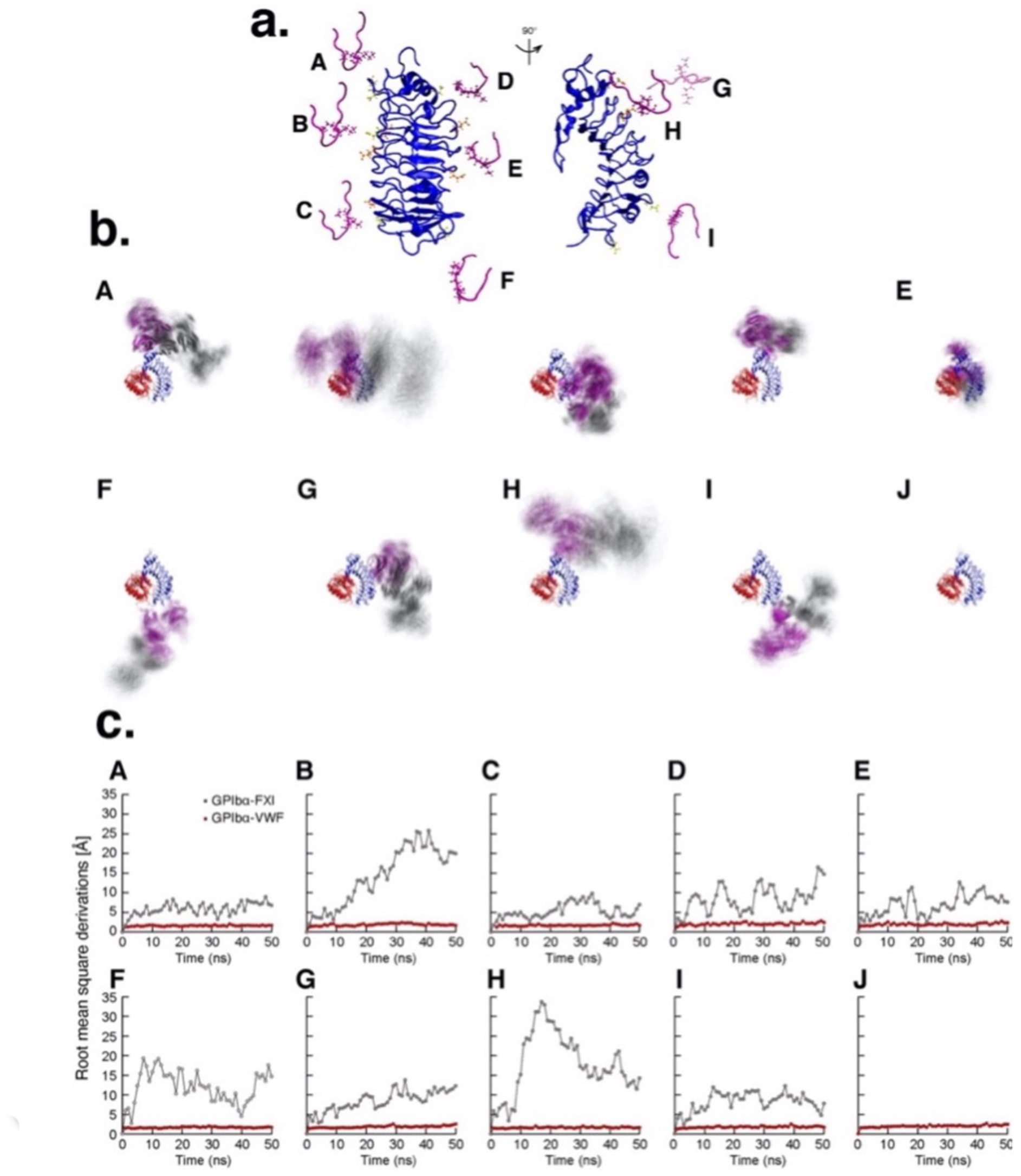
Structural Fluctuation of VWF-GPIbα and FXI-GPIbα. **(a)** Nine different initial positions of FXI relative to the GPIbα tested are shown. These initial position of FXI was determined based on the negatively charged amino acids depicted as yellow (aspartic acid: D) and orange (glutamic acid:E) on GPIbα. The left and right images show the views of GPIbα from the opposite side of the VWF binding site, and the views after a 90-degree clockwise rotation around the vertical axis from that position**. (b)** The time-lapse images of VWF(red)-GPIbα (blue) binding structures in the presence of FXI (purple) at 9 different initial positions described in panel **a**, and a control experiment in the absence of FXI. The snapshots were overlayed every 1 ns between 400 ns and 450 ns with an alpha value of 0.02. The results are also shown as **Supplemental Movies 2. (c)** The root mean square deviations (RMSDs) of atoms constructing VWF-GPIbα (red) and FXI-GPIbα (gray), excluding the water molecules, for 400 to 450 ns are shown. The structure at 400 ns was used as reference. Sub-panel A-I show the results started from the initial position of FXI corresponding to the same letter in panel **a**. Sub-panel J show the result in the absence of FXI.

## 3. The Root Mean Square Deviations (RMSDs) of Atoms

The RMSDs of all atoms, excluding the water molecules, were calculated every 10 ns from time 0 to 450 ns as a marker for the stability of the structural fluctuations of GPIbα-VWF and GPIbα-FXI. ^38^ The RMSDs calculated from time 0 to 450 ns are shown in **Supplemental Figure 2**. The calculation was considered stabilized when the continuous trend of increase in RMSDs disappeared. Based on the time-dependent changes in RMSD, the model stabilized after 250 ns in all initial starting positions of FXI tested. To compare the RMSDs after stabilization of the simulation, results within 400-450 ns were used to calculate RMSD for the structural stabilities of VWF-GPIbα and FXI-GPIbα. The average and standard deviations of the RMSD from 400 to 450 ns in all tested conditions are shown in **Supplemental Table 1**.

## 4. Non-Covalent Binding Energy Generated in VWF-GPIbα-VWF and FXI-GPIbα

The non-covalent binding energies in VWF-GPIbα and FXI-GPIbα bonds were calculated as an accumulation of the potential energy between amino acids forming salt badges ^39^ with the use of VMD and NAMD energy plugin (version 1.4). ^40, 41^ The salt bridges are constructed from two non-covalent interactions of hydrogen bond and ionic interactions. The non-covalent binding energy generated in VWF-GPIbα and FXI-GPIbα bonds in the entire calculation period are shown in **Supplemental Fig. 3.** To compare the non-covalent binding energy generated in VWF-GPIbα and FXI-GPIbα bonds after stabilization of the simulation calculation, results within 400-450 ns were compared as shown in **Fig. 3**. The means and standard deviations of calculated results from 400 to 450 ns are shown in **Supplemental Table 2**.

**Fig. 3.**
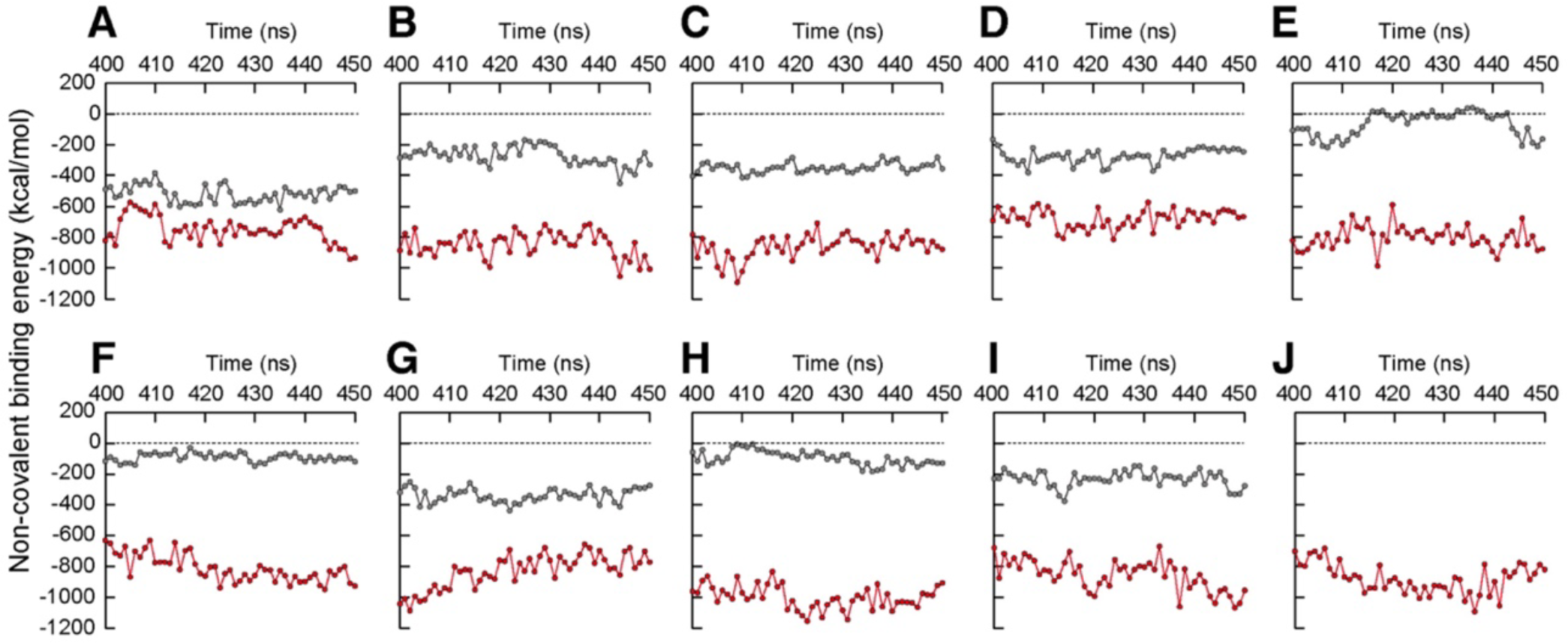
Non-Covalent Binding Energy for GPIbα-VWF and GPIbα-FXI binding. Time-dependent changes in the non-covalent binding energy generated between GPIbα binding with VWF in the presence of FXI at various initial positions (panel A to I) as well as its absence (panel J) from 400 ns to 450 ns are shown as red dots. The gray dots in panels A to I showed the non-covalent binding energy generated in GPIbα-FXI. The A-I corresponds to the initial position of FXI presented in the same letter in Fig 1A.

## 5. Salt Bridge Formation

The salt bridges were determined by the Salt Bridges Plugin (Version 1.1) as previously published. ^36, 39^ Salt bridges were considered to be formed when the distances between the anionic carboxylate (RCOO^−^) of either aspartic acid (ASP: N) or glutamic acid (GLU: E) and the cationic ammonium (RNH^3+^) from lysine (LYS: K) or the guanidinium (RNHC(NH_2_)^2+^) of arginine (ARG: R) were less than 4 Å in the 3-dimensional structure within GPIbα, VWF, and FXI. For GPIbα binding with VWF, there were 20 pairs of amino acids forming salt bridges. The percentages of the times the bridges were formed are shown as a heat map in **Fig. 4**. The values of the percentages of salt bridge formation from 400 to 450 ns between various amino acids starting from 9 initial positions of FXI and in the absence of FXI are shown in **Fig. 4**.

**Fig.4.**
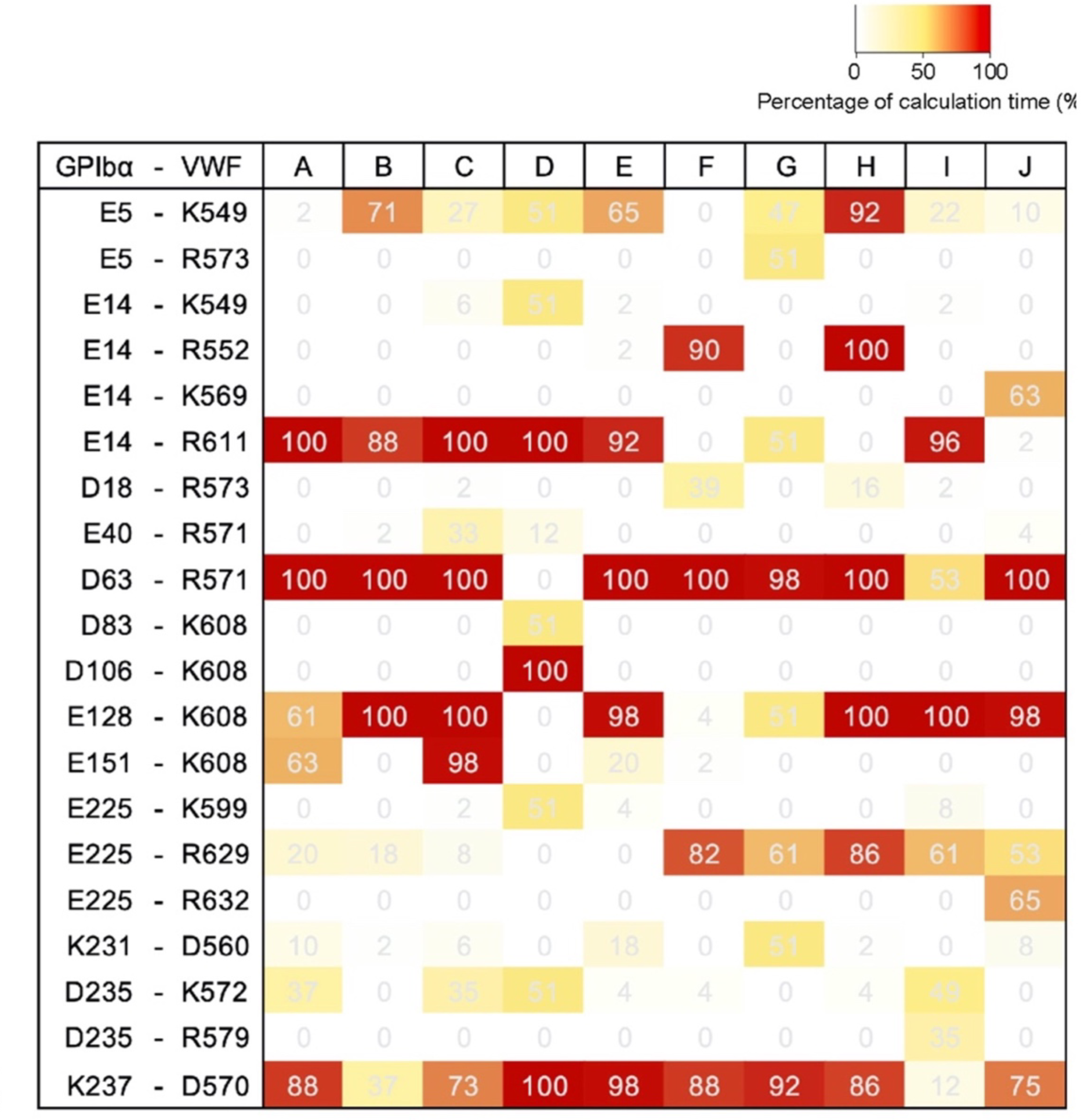
The Percentage of Time Where the Pairs of Amino Acids in GPIbα-VWF Formed Salt Bridges in the Presence and Absence of FXI Binding with GPIbα. The percentage of time the salt bridge was formed in each amino-acid pair in different initial FXI positions (A-I) and the absence of FXI (J). Results are shown as percentages within the simulation period of 400 to 450 ns. The A-I corresponds to the initial position of FXI presented in the same letter in Fig 1A. Panel J represents the results in the absence of FXI.

## Statistical analysis

Differences between RMSDs in atoms excluding water molecules, non-covalent binding energy between GPIbα-VWF and GPIbα-FXI were analyzed using 2-tailed Student’s t-tests. P values less than 0.05 were considered statistically significant.

## Data availability

The data that support the findings of this study are available from the corresponding. author upon reasonable request.

## Results

### 1. Difference in the Positional Fluctuations of Atoms Constructing VWF-GPIbα and FXI-GPIbα

Panel **a** of **Fig. 2** shows the 9 initial positions of FXI arranged around GPIbα. Panel **b** of **Fig. 2** shows the overlayed images of the snapshots obtained in every 1 ns from 400 to 450 ns of calculation. The original sequences of images obtained from 400 to 450 ns in every 1 ns of calculation are shown in **Supplemental Movie 1**. The structures of VWF-GPIbα were apparently more stable as compared to the FXI-GPIbα. The RMSDs of atoms constructing VWF-GPIbα bonds started from 9 different initial positions of FXI shown as **A** to **I** in panel **a** and **b** of **Fig. 2** were distributed from 1.6±0.1 to 2.1±0.3 Å. (**Table 1**) The RMSDs of atoms constructing FXI-GPIbα were distributed from 5.9±1.5 to 18.1±7.9 Å. (**Table 1**) The RMSDs in VWF-GPIbα were always smaller than those in FXI-GPIbα in all conditions as shown in **Table 1**. The time-dependent changes in RMSDs of atoms constructing VWF-GPIbα (red) and those constructing FXI-GPIbα (gray) within the last 50 ns of the whole calculation of 450 ns are shown in panel c of **Fig. 2**. **Supplemental Fig. 1** provides the time-dependent changes in RMSDs in atoms constructing VWF-GPIbα (red) and those constructing FXI-GPIbα (gray) from the beginning to the end of whole calculation. **Supplemental movie 2** provides all calculation results from the beginning to the end of 450 ns of calculations.

**Table 1.**
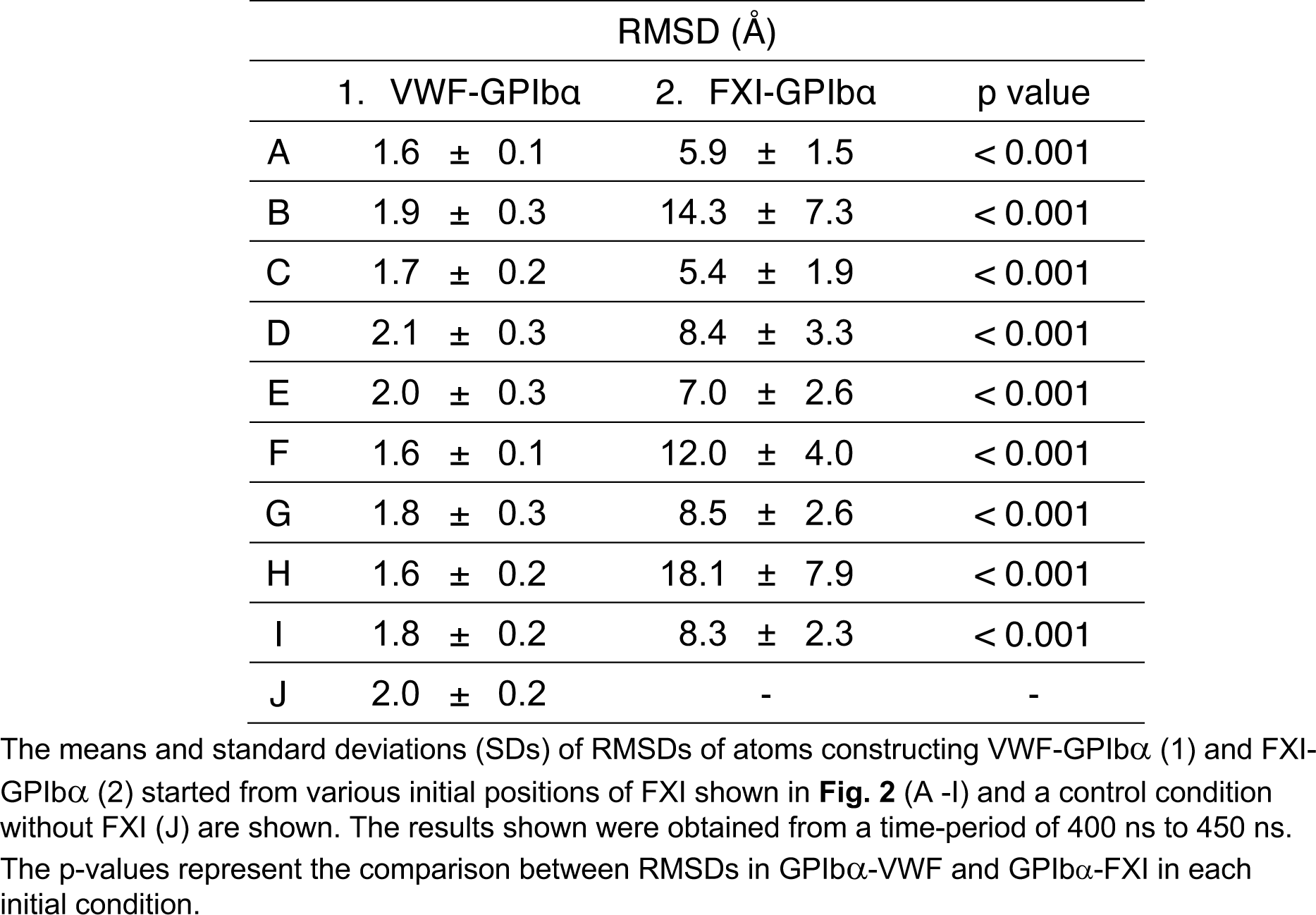
The Means and Standard Deviations of RMSDs of Atoms Constructing VWF-GPIbα and FXI-GPIbα from 400 ns to 450 ns.

### 2. Non-Covalent Binding Energy Generated in VWF-GPIbα and FXI-GPIbα

The non-covalent binding energy generated in VWF-GPIbα calculated from 9 initial conditions were distributed from −678.5±58.3 to −1000.4±75.1 kcal/mol. (**Table 2**) Those generated in FXI-GPIbα were distributed from −65.5±79.7 to −517.6±54.2 kcal/mol, and the values were lower as compared to the value in VWF-GPIbα. (**Table 2) Fig 3** show the time-dependent changes in the non-covalent binding energy generated in VWF-GPIbα (red) and FXI-GPIbα (gray) at the last 50 ns of the whole 450 ns of calculations. The non-covalent binding energy generated in VWF-GPIbα in the absence of FXI of −879.7 ± 96.3 kcal/mol was not markedly different from the values in its absence. (**Table 2**) **Supplemental Fig. 2** provides the time-dependent changes in non-covalent binding energy generated in VWF-GPIbα bond (red) and in the FXI-GPIbα bond (gray) from the beginning to the end of the whole calculation.

**Table 2.** The Mean and Standard Deviations of Non-Covalent Binding Energy Generated between GPIbα-VWF and GPIbα-FXI.

| Non-covalent Binding energy (kcal/mol) |  |  |  |
| --- | --- | --- | --- |
| | GPIIbα-VWF | GPIIbα-FXI | p value |
| A | -750.4 ± 86.3 | -517.6 ± 54.2 | < 0.001 |
| B | -845.3 ± 81.0 | -271.2 ± 60.0 | < 0.001 |
| C | -860.1 ± 75.3 | -348.1 ± 31.3 | < 0.001 |
| D | -678.5 ± 58.3 | -270.8 ± 46.1 | < 0.001 |
| E | -799.7 ± 73.5 | -65.5 ± 79.7 | < 0.001 |
| F | -812.7 ± 87.8 | -91.2 ± 27.2 | < 0.001 |
| G | -829.9 ± 109.0 | -337.1 ± 44.9 | < 0.001 |
| H | -1000.4 ± 75.1 | -95.5 ± 45.7 | < 0.001 |
| I | -857.2 ± 100.3 | -231.4 ± 50.0 | < 0.001 |
| J | -879.7 ± 96.3 | - | - |

### 3. Salt Bridges Formation between GPIbα and VWF

The pairs of amino acids forming salt bridges for more than 50% of calculation periods were colored as shown in the heat map in **Fig. 4**. There were 5 pairs of salt bridges in the absence of FXI. (panel **J**, **Fig. 4**) The numbers of the pairs of salt bridges formed more than 50% of calculation period after 400 ns in the presence of FXI varies from 3 (panel **D**, **Fig. 4**) to 6 (panel **H**, **Fig. 4**).

## Discussion

The biological functions and physical parameters of the platelet membrane protein GPIbα binding with VWF (VWF-GPIbα bond) and with FXI (FXI-GPIbα bond) differ substantially. (**Fig. 5**) The VWF-GPIbα bonds support platelet adhesion resistant to the detaching force generated by blood flow. The positional fluctuations of atoms constructing the VWF-GPIbα bond were smaller than that of the FXI-GPIbα bond in all 9 different initial conditions tested. The amount of non-covalent binding energy generated in VWF-GPIbα was larger than in FXI-GPIbα in all conditions tested. The probabilities of the pair of amino acids forming salt bridges in the VWF-GPIbα bond were always higher than the probabilities for the FXI-GPIbα bond. Even though its larger size, FXI binding with GPIbα minimally influenced the structures and functions of the VWF-GPIbα bond.

**Fig. 5.**
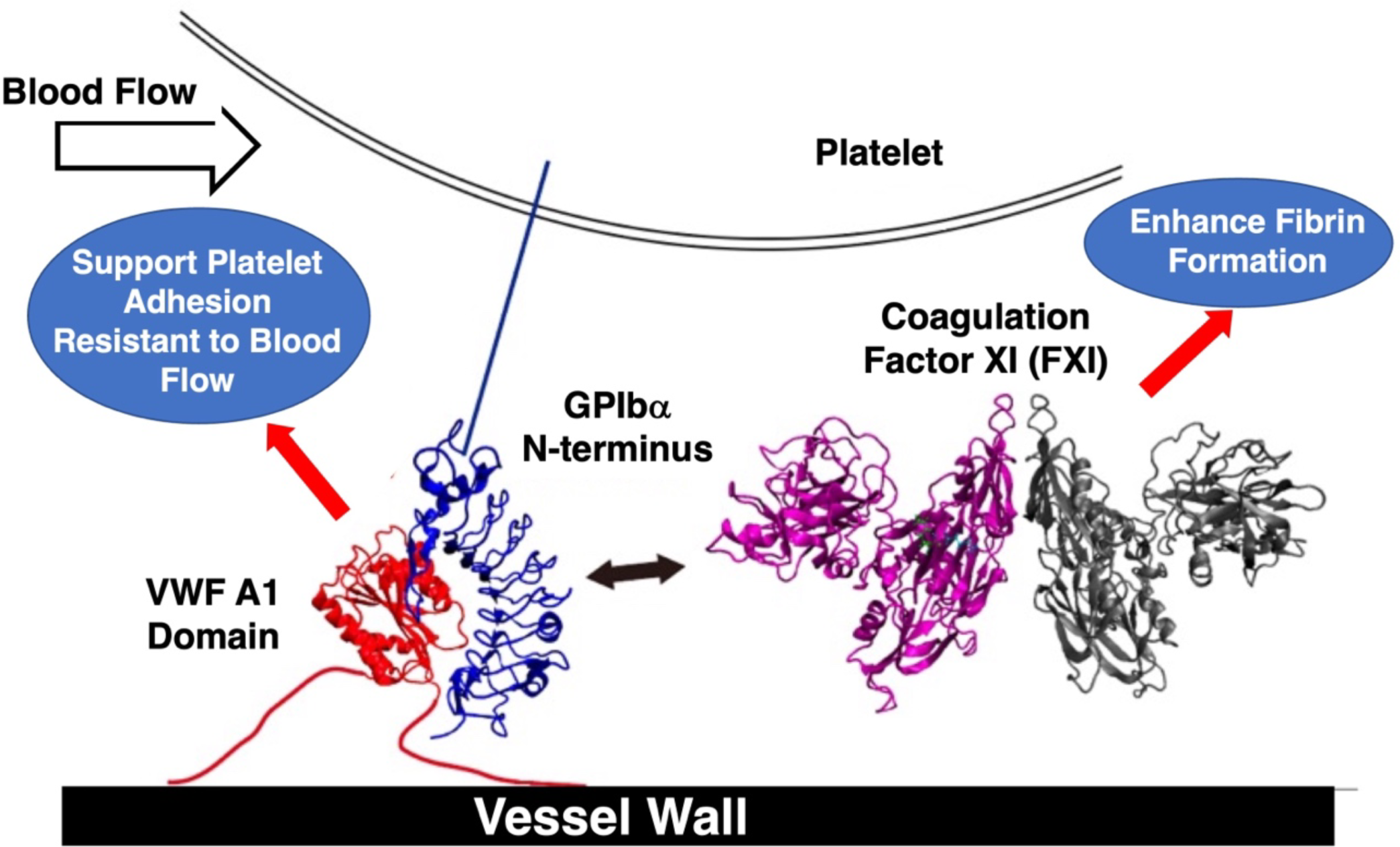
Distinct Biological Roles of Platelet Membrane Glycoprotein Ibα on Platelet Adhesion Mediated by VWF Binding and on Coagulation Mediated by FXI Binding. The platelet GPIbα molecules support platelet adhesion resistant to the detaching force generated by their binding with VWF. This biological function is characterized as a stable binding structure with higher non-covalent binding energy generated in the VWF-GPIbα bond. The GPIbα molecules also enhance coagulation by their binding with FXI. This biological function was mediated by unstable binding with less non-covalent binding energy generated in FXI-GPIbα.

MD simulation is not a novel technic, ^42^ but has become an important tool for clarifying the biological functions of various proteins due to rapid progress in the calculation power of high-performance computers. ^43–45^ The establishment of coarse-grained force fields based on quantum mechanics into molecular mechanics such as the Chemistry at HARvard Macromolecular Mechanics (CHARMM) force field^19, 46^ helped to generate comprehensive results from MD calculations.^42^ Due to theoretical limitations, the validity of the calculated results of MD simulation should be confirmed by other methodologies, such as biological experiments. For the interactions between VWF and GPIbα, the validity of the results of MD calculations was confirmed quantitatively by comparing the predicted binding force ^21^ with the biological measurement by atomic force microscopy^22^ and optical tweezer.^23^ The validity of the predicted binding structure of VWF-GPIbα was further established by predicting the phenotype of the platelet-type von Willebrand disease caused by the G233D mutant in GPIbα.^24^ The lower non-covalent binding energy in GPIbα-VWF bond was also shown in G233D mutant of GPIbα. ^25^ Our results showing the structural stability with more non-covalent binding energy generated in VWF-GPIbα bonds as compared to FXI-GPIbα shown here further support the specific physical characteristics of VWF-GPIbα mediating specific biological function.

The binding site of FXI for GPIbα has been reported as K252 and K253. ^26, 31^ However, the corresponding binding site(s) for FXI within GPIbα is yet to be clarified. Since K252 and K253 are positively charged amino acids, it is reasonable to assume that these amino acids start the interaction with negatively charged amino acids such as D and E on GPIbα. Thus, the MD simulations were started from 9 different positions of FXI binding with the locations of these negatively charged amino acids on GPIbα. Our experimental results suggest substantially greater structural fluctuation with less non-covalent binding energy in the FXI-GPIbα bond as compared to that in VWF-GPIbα. The minimal impacts of FXI binding with GPIbα on both the structural fluctuation and binding energies in GPIbα binding with VWF were also shown. FXI binding with GPIbα is neither stable nor generates strong non-covalent binding energies but may still be able to enhance local coagulation activity around platelet by increasing the probability of FXI interaction with thrombin. ^26^ ^47^ The two distinct physical characteristics in the VWF-GPIbα bond and the one in FXI-GPIbα shown here correspond to their specific biological functions.

The binding force generated by one pair of VWF-GPIbα was predicted and measured as approximately 50–100 pN. ^21–23^. Despite more than 10,000 molecules of GPIbα expressed on a single platelet cell,^18^ only a few pairs of GPIbα binding with VWF generate enough force to adhere platelet resisting to the blood flow. ^48^ Our current results suggest that at least five pairs of salt bridges were formed in VWF-GPIbα bonds with a probability higher than 50%. This means single platelet adhesion on VWF at a wall shear rate of 1,500 s^-1^ was supported by approximately 50 pairs of salt bridges formed longer than 50% of the calculation period. The importance of the roles of amino acids forming salt bridge in VWF-GPIbα bond previously reported as D570,^49^ K572,^50^ R571,^50^ ^51^ R611,^50, 52^ in VWF and K237,^53^ D235,^54^ E128,^55^ and E14,^55^ in GPIbα were confirmed in our experiments.

Our study had several methodological limitations. First, the behavior of atoms sharing electric clouds cannot be determined theoretically. ^56^ The CHARMM force field that we used in this analysis is an established methodology to deal with simulation requiring the consideration of quantum mechanics, but coarse-grained quantum mechanics in the CHARMM force field might cause potential errors in our MD calculations. Moreover, Poincaré JH theoretically demonstrated that three-body problems, in general, cannot be solved analytically. ^57^ Apparently, our MD simulation targeted three proteins (GPIbα, VWF, and FXI). To obtain robust results, our MD calculations were started from 9 different initial positions of FXI. Since the initial structures were determined only based on the electrical charge of amino acid and not from results of established biological methods (such as X-ray crystallography), some of the initial structures we used in our experiments may not have biological relevance. However, our results supporting the 2 distinct characteristics of less structural fluctuation with greater non-covalent binding energy in GPIbα-VWF as compared to GPIbα-FXI is valid given the robustness of the findings across various possible initial positions of FXI. Thus, despite limitations, our conclusions support two distinct binding characteristics of platelet GPIbα bonds: the one that mediates platelet adhesion on VWF requires a strong binding force; and the other mediating procoagulant activity of platelets by localization of FXI, should not be influenced.

In conclusion, MD simulation revealed structural instability and smaller non-covalent binding energy in the FXI-GPIbα bond as compared to the one in VWF-GPIbα. These structural characteristics of VWF-GPIbα and FXI-GPIbα are in agreement with the 2 distinct mechanisms of molecular functions: one producing strong enough binding energy to support platelet adhesion resistant to the blood flow; the other helping enhanced coagulation by increasing the probability of FXI interaction with thrombin around platelets. MD simulation may provide physical insight into various biological functions of protein bonds.

## Acknowledgement

We acknowledge the support by Strategic Program for Innovational Research Field 1 for Super-computational Life Science, grant-in-aid for MEXT/JSPS KAKENHI 19H03661, AMED grant number A368TS, Bristol-Myers Squibb for independent research support project (33999603) and a grant from Nakatani Foundation for Advancement of Measuring Technologies in Biomedical Engineering and Vehicle Racing Commemorative Foundation (6236).

## Supplementary Materials

**Supplemental Fig. 1. The Root Mean Square Derivations of Atoms Constructing VWF-GPIbα and FXI-GPIbα during the Entire Calculation Period.**

The root mean square deviations (RMSDs) of atoms, excluding the water molecules, constructing GPIbα-VWF (red) and GPIbα-FXI calculated for 450 ns in every 10 ns. The A-I corresponds to the initial position of FXI presented in the same letter in Fig 1A. Panel J represents the results in the absence of FXI.

**Supplemental Fig. 2. The Non-Covalent Binding Energy for GPIbα-VWF and GPIbα-FXI during Entire Calculation Period.**

The non-covalent binding energy in GPIbα-VWF (red) and GPIbα-FXI (black) calculated from time 0 to 450 ns in every 10 ns are shown. The A-I corresponds to the initial position of FXI preseted in the same letter in Fig 1A. The panel J represent the results in the absence of FXI.

**Supplemental Movie 1. The Results of Molecular Dynamic Calculation of GPIbα, VWF, and FXI in Conditions with Various Initial Positions of FXI from 400 ns to 450 ns.**

The Supplemental Movie 1 file is provided separately from the manuscript file with the file name of Supplemental Movie1.mp4. Panels A to I show the results of MD calculation expressed as the sequential snapshot images obtained every 1 ns from time 400 to time 450 ns in the presence of FXI arranged at various sites on GPIbα. Panel I show the same calculation results but in the absence of FXI. The movies were constructed from the position coordinates of atoms constructing GPIbα, and VWF, in the presence and absence of FXI at the rate of 10 fps.

**Supplemental Movie 2. The Entire Results of Molecular Dynamic Calculation of GPIbα, VWF and FXI in Conditions with Various Initial Positions of FXI.**

The Supplemental Movie 2 file is provided separately from the manuscript file with the file name of Supplemental Movie2.mp4. Panels A to I show the results of MD calculation expressed as the sequential snapshot images obtained every 10 ns from time 0 to time 450 ns in the presence of FXI arranged at various sites on GPIbα. Panel I show the same calculation results but in the absence of FXI. The movies were constructed from the position coordinates of atoms constructing GPIbα, and VWF, in the presence and absence of FXI at the rate of 10 fps.

Supplemental pdb files (A.pdb to I.pdb) Initial Structure of GPIbα, VWF, and FXI at Various Conditions Expressed as pdb File.

The pdb file is provided separately from the manuscript file. Each pdb file is provided as a.pdb to i.pdb in supplemental pdb files. All pdb files provide here represent the initial structure of GPIbα, VWF, and FXI in our experiments.

## Reference

1. Ruggeri ZM. Platelets in atherothrombosis. Nature medicine. 2002;8:1227–1234.

2. Goto S, Hasebe T, Takagi S. Platelets: Small in Size But Essential in the Regulation of Vascular Homeostasis - Translation From Basic Science to Clinical Medicine. Circulation journal: official journal of the Japanese Circulation Society. 2015;79:1871–1881. doi: 10.1253/circj.CJ-14-1434

3. Hagihara M, Higuchi A, Tamura N, Ueda Y, Hirabayashi K, Ikeda Y, Kato S, Sakamoto S, Hotta T, Handa S. Platelets, after exposure to a high shear stress, induce IL-10-producing, mature dendritic cells in vitro. The Journal of Immunology. 2004;172:5297–5303.

4. Koupenova M, Ravid K. Biology of platelet purinergic receptors and implications for platelet heterogeneity. Frontiers in pharmacology. 2018;9:37.

5. Kawamura Y, Takahari Y, Tamura N, Eguchi Y, Urano T, Ishida H, Goto S. Imaging of structural changes in endothelial cells and thrombus formation at the site of FeCl3-induced injuries in mice cremasteric arteries. Journal of atherosclerosis and thrombosis. 2010;16:807–814.

6. Tamura N, Kitajima I, Kawamura Y, Toda E, Eguchi Y, Ishida H, Goto S. Important regulatory role of activated platelet-derived procoagulant activity in the propagation of thrombi formed under arterial blood flow conditions. Circulation Journal. 2009;73:540–548.

7. Savage B, Almus-Jacobs F, Ruggeri ZM. Specific synergy of multiple substrate-receptor interactions in platelet thrombus formation under flow. Cell. 1998;94:657–666.

8. Savage B, Saldívar E, Ruggeri ZM. Initiation of platelet adhesion by arrest onto fibrinogen or translocation on von Willebrand factor. Cell. 1996;84:289–297.

9. Miyata S, Goto S, Federici AB, Ware J, Ruggeri ZM. Conformational changes in the A1 domain of von Willebrand factor modulating the interaction with platelet glycoprotein Ibalpha. The Journal of biological chemistry. 1996;271:9046–9053.

10. Madabhushi SR, Zhang C, Kelkar A, Dayananda KM, Neelamegham S. Platelet GpIbα Binding to von Willebrand Factor Under Fluid Shear: Contributions of the D’D3-Domain, A1-Domain Flanking Peptide and O-Linked Glycans. Journal of the American Heart Association. 2014;3:e001420.

11. Goto S, Ikeda Y, Saldivar E, Ruggeri ZM. Distinct mechanisms of platelet aggregation as a consequence of different shearing flow conditions. The Journal of clinical investigation. 1998;101:479–486. doi: 10.1172/JCI973

12. Coller BS, Peerschke EI, Scudder LE, Sullivan CA. Studies with a murine monoclonal antibody that abolishes ristocetin-induced binding of von Willebrand factor to platelets: additional evidence in support of GPIb as a platelet receptor for von Willebrand factor. 1983.

13. Girma J, Takahashi Y, Yoshioka A, Diaz J, Meyer D. Ristocetin and botrocetin involve two distinct domains of von Willebrand factor for binding to platelet membrane glycoprotein lb. Thrombosis and haemostasis. 1990;64:326–332.

14. Ruggeri ZM, Pareti FI, Mannucci PM, Ciavarella N, Zimmerman TS. Heightened interaction between platelets and factor VIII/von Willebrand factor in a new subtype of von Willebrand’s disease. New England Journal of Medicine. 1980;302:1047–1051.

15. De Marco L, Mazzucato M, De Roia D, Casonato A, Federici AB, Girolami A, Ruggeri ZM. Distinct abnormalities in the interaction of purified types IIA and IIB von Willebrand factor with the two platelet binding sites, glycoprotein complexes Ib-IX and IIb-IIIa. The Journal of clinical investigation. 1990;86:785–792.

16. Ginsburg D, Sadler EJ. Von Willebrand disease: a database of point mutations, insertions, and deletions. Thrombosis and haemostasis. 1993;69:177–184.

17. Budde U, Schneppenheim R. von Willebrand Factorandvon Willebrand Disease. Reviews in clinical and experimental hematology. 2001;5:335–335.

18. Goto S, Salomon DR, Ikeda Y, Ruggeri ZM. Characterization of the unique mechanism mediating the shear-dependent binding of soluble von Willebrand factor to platelets. The Journal of biological chemistry. 1995;270:23352–23361.

19. Brooks BR, Brooks CL, 3rd, Mackerell AD, Jr., Nilsson L, Petrella RJ, Roux B, Won Y, Archontis G, Bartels C, Boresch S, et al. CHARMM: the biomolecular simulation program. Journal of computational chemistry. 2009;30:1545–1614. doi: 10.1002/jcc.21287

20. Brooks BR, Bruccoleri RE, Olafson BD, States DJ, Swaminathan S, Karplus M. CHARMM: A program for macromolecular energy, minimization, and dynamics calculations. Journal of computational chemistry. 1983;4:187–217. doi: https://doi.org/10.1002/jcc.540040211

21. Shiozaki S, Takagi S, Goto S. Prediction of Molecular Interaction between Platelet Glycoprotein Ibα and von Willebrand Factor using Molecular Dynamics Simulations. Journal of atherosclerosis and thrombosis. 2016;23:455–464. doi: 10.5551/jat.32458

22. Tobimatsu H, Nishibuchi Y, Sudo R, Goto S, Tanishita K. Adhesive Forces between A1 Domain of von Willebrand Factor and N-terminus Domain of Glycoprotein Ibα Measured by Atomic Force Microscopy. Journal of atherosclerosis and thrombosis. 2015;22:1091–1099. doi: 10.5551/jat.28423

23. Kim J, Zhang CZ, Zhang X, Springer TA. A mechanically stabilized receptor-ligand flex-bond important in the vasculature. Nature. 2010;466:992–995. doi: 10.1038/nature09295

24. Goto S, Oka H, Ayabe K, Yabushita H, Nakayama M, Hasebe T, Yokota H, Takagi S, Sano M, Tomita A, et al. Prediction of binding characteristics between von Willebrand factor and platelet glycoprotein Ibα with various mutations by molecular dynamic simulation. Thromb Res. 2019;184:129–135. doi: 10.1016/j.thromres.2019.10.022

25. Nakayama M, Goto S, Goto S. Physical Characteristics of von Willebrand Factor Binding with Platelet Glycoprotein Ibɑ Mutants at Residue 233 Causing Various Biological Functions. TH Open. 2022;6:e421–e428.

26. Baglia FA, Badellino KO, Li CQ, Lopez JA, Walsh PN. Factor XI binding to the platelet glycoprotein Ib-IX-V complex promotes factor XI activation by thrombin. The Journal of biological chemistry. 2002;277:1662–1668. doi: 10.1074/jbc.M108319200

27. Baglia FA, Gailani D, López JA, Walsh PN. Identification of a binding site for glycoprotein Ibα in the Apple 3 domain of factor XI. Journal of Biological Chemistry. 2004;279:45470–45476.

28. Dumas JJ, Kumar R, Seehra J, Somers WS, Mosyak L. Crystal Structure of the GpIbα-Thrombin Complex Essential for Platelet Aggregation. Science (New York, NY). 2003;301:222–226. doi: 10.1126/science.1083917

29. Kossmann S, Lagrange J, Jäckel S, Jurk K, Ehlken M, Schönfelder T, Weihert Y, Knorr M, Brandt M, Xia N, et al. Platelet-localized FXI promotes a vascular coagulation-inflammatory circuit in arterial hypertension. Sci Transl Med. 2017;9. doi: 10.1126/scitranslmed.aah4923

30. Papagrigoriou E, McEwan PA, Walsh PN, Emsley J. Crystal structure of the factor XI zymogen reveals a pathway for transactivation. Nature structural & molecular biology. 2006;13:557–558.

31. Baglia FA, Shrimpton CN, Emsley J, Kitagawa K, Ruggeri ZM, López JA, Walsh PN. Factor XI interacts with the leucine-rich repeats of glycoprotein Ibα on the activated platelet. Journal of Biological Chemistry. 2004;279:49323–49329.

32. Phillips JC, Braun R, Wang W, Gumbart J, Tajkhorshid E, Villa E, Chipot C, Skeel RD, Kalé L, Schulten K. Scalable molecular dynamics with NAMD. Journal of computational chemistry. 2005;26:1781–1802. doi: 10.1002/jcc.20289

33. Boonstra S, Onck PR, van der Giessen E. CHARMM TIP3P water model suppresses peptide folding by solvating the unfolded state. The journal of physical chemistry B. 2016;120:3692–3698.

34. Li P, Roberts BP, Chakravorty DK, Merz KM, Jr. Rational Design of Particle Mesh Ewald Compatible Lennard-Jones Parameters for +2 Metal Cations in Explicit Solvent. Journal of chemical theory and computation. 2013;9:2733–2748. doi: 10.1021/ct400146w

35. Abraham MJ, Gready JE. Optimization of parameters for molecular dynamics simulation using smooth particle-mesh Ewald in GROMACS 4.5. Journal of computational chemistry. 2011;32:2031–2040.

36. Humphrey W, Dalke A, Schulten K. VMD: Visual molecular dynamics. Journal of molecular graphics. 1996;14:33–38. doi: https://doi.org/10.1016/0263-7855(96)00018-5

37. Goto S, Tamura N, Handa S, Arai M, Kodama K, Takayama H. Involvement of glycoprotein VI in platelet thrombus formation on both collagen and von Willebrand factor surfaces under flow conditions. Circulation. 2002;106:266–272. doi: 10.1161/01.cir.0000021427.87256.7e

38. Maiorov VN, Crippen GM. Significance of Root-Mean-Square Deviation in Comparing Three-dimensional Structures of Globular Proteins. Journal of Molecular Biology. 1994;235:625–634. doi: https://doi.org/10.1006/jmbi.1994.1017

39. Lee M, Chang HJ, Park JY, Shin J, Park JW, Choi JW, Kim JI, Na S. Conformational changes of Aβ (1–42) monomers in different solvents. Journal of Molecular Graphics and Modelling. 2016;65:8–14.

40. Ramadoss V, Dehez F, Chipot C. AlaScan: A graphical user interface for alanine scanning free-energy calculations. In: ACS Publications; 2016.

41. Buehl M, Wipff G. Insights into uranyl chemistry from molecular dynamics simulations. ChemPhysChem. 2011;12:3095–3105.

42. Alder BJ, Wainwright TE. Studies in Molecular Dynamics. I. General Method. The Journal of Chemical Physics. 1959;31:459–466. doi: 10.1063/1.1730376

43. Schreiber LR, Bluhm H. Toward a silicon-based quantum computer. Science (New York, NY). 2018;359:393–394. doi: 10.1126/science.aar6209

44. Sravanthi G, Grace B, Kamakshamma V. A review of High Performance Computing. IOSR Journal of Computer Engineering. 2014;16:36–43.

45. Goto S, McGuire DK, Goto S. The Future Role of High-Performance Computing in Cardiovascular Medicine and Science-Impact of Multi-Dimensional Data Analysis. Journal of atherosclerosis and thrombosis. 2021;advpub. doi: 10.5551/jat.RV17062

46. Huang J, MacKerell AD, Jr. CHARMM36 all-atom additive protein force field: validation based on comparison to NMR data. Journal of computational chemistry. 2013;34:2135–2145. doi: 10.1002/jcc.23354

47. Ruggeri ZM, Zarpellon A, Roberts JR, Mc Clintock RA, Jing H, Mendolicchio GL. Unravelling the mechanism and significance of thrombin binding to platelet glycoprotein Ib. Thrombosis and haemostasis. 2010;104:894–902. doi: 10.1160/th10-09-0578

48. Tamura N, Shimizu K, Shiozaki S, Sugiyama K, Nakayama M, Goto S, Takagi S, Goto S. Important Regulatory Roles of Erythrocytes on Platelet Adhesion to the von Willebrand Factor on the Wall Under Blood Flow Conditions. Thrombosis and haemostasis. 2021.

49. Bonnefoy A, Yamamoto H, Thys C, Kito M, Vermylen J, Hoylaerts MF. Shielding the front-strand beta 3 of the von Willebrand factor A1 domain inhibits its binding to platelet glycoprotein Ibalpha. Blood. 2003;101:1375–1383. doi: 10.1182/blood-2002-06-1818

50. Matsushita T, Meyer D, Sadler JE. Localization of von willebrand factor-binding sites for platelet glycoprotein Ib and botrocetin by charged-to-alanine scanning mutagenesis. The Journal of biological chemistry. 2000;275:11044–11049. doi: 10.1074/jbc.275.15.11044

51. Shimizu A, Matsushita T, Kondo T, Inden Y, Kojima T, Saito H, Hirai M. Identification of the amino acid residues of the platelet glycoprotein Ib (GPIb) essential for the von Willebrand factor binding by clustered charged-to-alanine scanning mutagenesis. The Journal of biological chemistry. 2004;279:16285–16294. doi: 10.1074/jbc.M307230200

52. Castaman G, Eikenboom JC, Rodeghiero F, Briët E, Reitsma PH. A novel candidate mutation (Arg611-->His) in type I ’platelet discordant’ von Willebrand’s disease with desmopressin-induced thrombocytopenia. Br J Haematol. 1995;89:656–658. doi: 10.1111/j.1365-2141.1995.tb08383.x

53. Fontayne A, De Maeyer B, De Maeyer M, Yamashita M, Matsushita T, Deckmyn H. Paratope and epitope mapping of the antithrombotic antibody 6B4 in complex with platelet glycoprotein Ibalpha. The Journal of biological chemistry. 2007;282:23517–23524. doi: 10.1074/jbc.M701826200

54. Enayat S, Ravanbod S, Rassoulzadegan M, Jazebi M, Tarighat S, Ala F, Emsley J, Othman M. A novel D235Y mutation in the GP1BA gene enhances platelet interaction with von Willebrand factor in an Iranian family with platelet-type von Willebrand disease. Thrombosis and haemostasis. 2012;108:946–954. doi: 10.1160/th12-04-0189

55. Shen Y, Cranmer SL, Aprico A, Whisstock JC, Jackson SP, Berndt MC, Andrews RK. Leucine-rich repeats 2-4 (Leu60-Glu128) of platelet glycoprotein Ibalpha regulate shear-dependent cell adhesion to von Willebrand factor. The Journal of biological chemistry. 2006;281:26419–26423. doi: 10.1074/jbc.M604296200

56. Carnio EG, Breuer HP, Buchleitner A. Wave-Particle Duality in Complex Quantum Systems. J Phys Chem Lett. 2019;10:2121–2129. doi: 10.1021/acs.jpclett.9b00676

57. Chenciner A. Poincaré and the three-body problem. In: Henri Poincaré, 1912–2012. Springer; 2015:51–149.

